# The ontogeny of chimpanzee technological efficiency

**DOI:** 10.1101/2024.07.31.605100

**Authors:** Sophie Berdugo, Emma Cohen, Arran J. Davis, Tetsuro Matsuzawa, Susana Carvalho

**Author notes:** **Author Contributions: S.B.** conceived of the study, designed, and coordinated the study, collected data, analysed and visualised the data, and wrote the manuscript. **E.C.** provided supervision, participated in the design of the study, and commented on the manuscript. **A.J.D.** analysed and visualised the data and commented on the manuscript. **T.M.** collected the original data set and commented on the manuscript. **S.C.** collected the original data set, provided supervision, participated in the design of the study, and commented on the manuscript. **Competing Interest Statement:** The authors declare no conflict of interest.

## Abstract

Primate extractive foraging requires years of dedicated learning. Throughout this period, learners peer at conspecifics engaging in the behaviour (“models”), interacting with the model and their tools, and sometimes stealing the freshly extracted resource. This also corresponds to an extended period of tolerance from the models. Yet the long-term effect of variation in experiences during this period on the technological efficiency of individuals is unknown for primate tool use, and no research has assessed the role of both the learner and the model(s) in generating individual differences. Using >680 hours of video spanning 25 years, we assessed whether experiences during the stone tool use social learning period (“early learning period”; ages 0–5) predicted the post-early learning period (ages 6+) technological efficiency in wild chimpanzees in Bossou, Guinea. We found that learners varied in how frequently they peered at the models’ whole nut-cracking bouts, how many learning opportunities their mothers presented, and the amount of tolerance and intolerance they experienced from all selected models. Learners who experienced more intolerance became less efficient tool users, whereas learners who were exposed to more social learning opportunities and tolerance became more efficient. Peering at the whole nut-cracking bout decreased subsequent efficiency, hinting at learners acquiring less efficient cultural components of the behaviour. Our findings highlight the role of social learning in the acquisition of stone tool use and support the view that social learning opportunities within a tolerant environment are key in explaining the emergence and maintenance of complex forms of primate technology.

**Significance Statement:** The capacity and inclination to learn from others, along with social learning opportunities provided by tolerant groupmates, are thought to have enabled the evolution of technology in primates, including hominins. The influence of the learning period on long-term individual variation in technological efficiency remains unknown for non-human primates but has significant implications for cultural transmission and evolution. We provide longitudinal support for the hypothesis that exposure to social learning opportunities during development predicts subsequent technological efficiency. Moreover, we show that low amounts of *in*tolerance, not just general tolerance, is key in the ontogeny of technological efficiency. Finally, we find aspects of behavioural acquisition relating to accurate transmission of cultural traits rather than to learning to use tools efficiently.

## Introduction

Variation in tool use behaviour is found across wild primates, including in tufted capuchins (*Sapajus libidinosus*) (1), long-tailed macaques (*Macaca fascicularis*) (2, 3), Sumatran orangutans (*Pongo pygmaeus abelii*) (4), and chimpanzees (*Pan troglodytes* spp.) (5). Such variation is established at the community (6, 7), age- and sex-class (1, 8), and individual levels (9, 10). Variation in the efficiency (∝benefit/cost) (11), as opposed to simply the frequency, of extracting calorie-rich and nutrient-dense resources directly impacts energy expenditure (12), as well as being a driver in cultural and technological evolution (9, 11, 13).

Previous short-term studies on perishable tool use established that differences in behaviour stem from systematic variation in social learning experiences (4, 5, 14)—learning involving observing or interacting with another individual or their products (15). However, the ontogenetic causes of individual differences are unknown for stone tools, and the long-term consequences of divergent learning experiences have never been explored in primate technology. Investigating how and why this variation in technological efficiency emerges will inform our understanding of processes with profound evolutionary consequences (9, 16), including potential individual variation in social learning^13^ and theories of the origins of tool use (17).

For culture to evolve, the cultural trait and its variants must be inherited by the subsequent generation (18). As such, cultural information transfer must be vertical (from older kin) or oblique (from older non-kin), with solely horizontal transmission (peer-to-peer) being inadequate (18). Therefore, along with necessary ecological and manual dexterity prerequisites, exposure to suitable social learning opportunities enabled primates, including hominins, to evolve material culture (17). Van Schaik and colleagues argued that opportunities to learn complex skills socially can only be exploited where the older competent conspecifics are also tolerant of the younger generation learning over prolonged periods (17). A mathematical model on tool-using skills provided preliminary support for this hypothesis by demonstrating that populations with sufficient social tolerance and opportunities for social learning are more likely to evolve complex skills, such as tool use (19).

Investigating the developmental drivers of individual variation in chimpanzee extractive-foraging techniques can also help advance our understanding of primate cognition and cognitive evolution. Carvajal & Schuppli argued that social learning opportunities are a central component of the evolution of complex cognition as they can dictate the ease and speed with which fitness-enhancing skills and information are obtained (20). The propensity to learn faithfully from others—which enabled the intergenerational accumulation of knowledge—propelled hominins into the “cultural niche”, which supported their ecological success (21). Given that our knowledge of early hominin technology is currently constrained to non-perishable stone tools, it is important to establish the processes generating variation in primate stone tool use to better capture the scope of hominin cognitive capabilities (22).

West African chimpanzees (*P. t. verus*) use stone and wooden hammers to crack nuts (23, 24), with the technique varying cross-culturally (6). The chimpanzees of Bossou, Guinea, use a stone hammer-and-anvil composite to extract oil palm nut kernels (*Elaeis guineensis*) (25), with individuals varying in their technological efficiency (9). The skill requires coordination of both hands whilst manoeuvring the three components (the hammer, anvil, and nut) (26), and is developed over a prolonged early learning period (hereafter, “ELP”) proposed to last 3–5 years (8, 27). The successful sequencing of the fundamental nut-cracking actions (take nut, place nut, hold stone, hit nut, and eat kernel) becomes more formalised with age, with sequences consisting of four of the five actions only emerging from 3.5 years old (27). This highlights the complexity of the behaviour and the importance of sustaining attention to observe the entire nut-cracking bout of knowledgeable models.

Nut-cracking is acquired via a combination of individual and social learning (7) during an “education by master-apprenticeship” learning process, which necessitates a tolerant model (the “master,” typically the mother) for the learner (the “apprentice”; see Figure 1) (28). Two mothers in Bossou (Nina and Pama) could not crack nuts—they are thought to have emigrated from a community without the behaviour—and yet all of their infants (Na, Nto, Pili, Poni, and Peley) learned the skill (23). This illustrates the importance of non-maternal models acting as “masters” for the “apprentices”.

**Figure 1.**
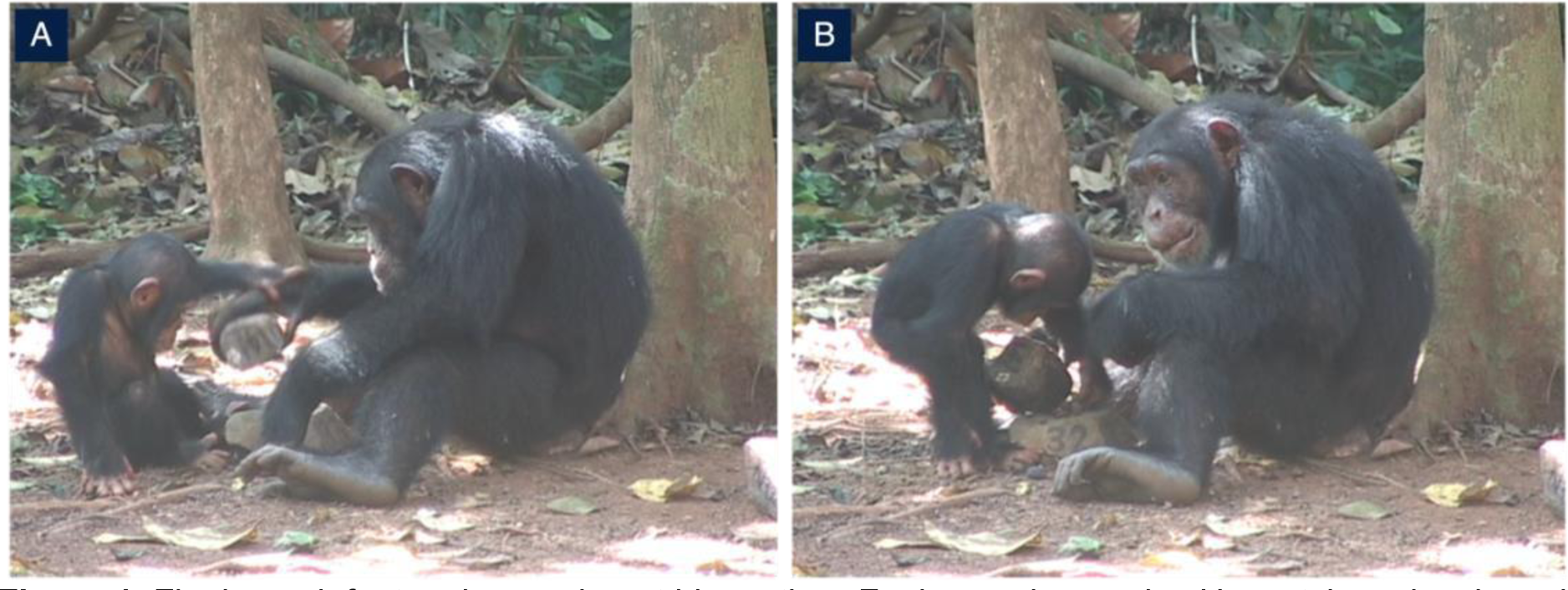
Flanle, an infant male, peering at his mother, Fanle, as she cracks. He watches closely and rests his hand on Fanle’s to learn the energy required to crack oil palm nuts (A). He then scrounges the extracted kernel (B). Image credit: Tetsuro Matsuzawa.

Nut-cracking with a stone hammer-and-anvil composite cannot be individually re-innovated in the wild— a competent conspecific is required to demonstrate the nut-cracking technique and motivate the learner to engage in the behaviour (29). Allowing infants to scrounge (i.e., take the extracted kernel to eat themselves) helps to maintain motivation to learn the behaviour over the prolonged period (30–32), though it has been suggested that the main impetus to learn the skill is to engage in the same behaviours as groupmates (28). Individuals may vary in their reliance on social learning, however, and the mechanisms used may vary systematically across individuals due to an ‘individual learning of social learning’ (13, 33, p. 215). As such, variation in both learner and model behaviour during the nut-cracking ELP is likely to influence the post-early learning period (hereafter “post-ELP”) tool use efficiency.

It is critical, therefore, to assess the combined influence of the learner and model(s) during the acquisition of complex skills at the individual level. Despite this, previous efforts have typically focused on establishing the source of sex differences, rather than of *individual* differences, in tool use, or have assessed the influence of singular components of the developmental period, such as the amount of time peering (i.e., ‘the attentive and sustained close-range watching of the activities of a conspecific’ (34, p. 5)) at the skill being performed. No prior research has assessed the role of both the learner and the model(s) in generating individual variation in technological behaviour.

Current hypotheses explaining the source of variation in technological behaviour are focused on explaining sex differences. Sex differences in tool-use are thought to emerge in infancy (35). An investigation into sex differences in “preparation” behaviours (i.e., engagement with sticks and woody vegetation) for ant-dipping by chimpanzees in Kalinzu, Uganda, found that although males manipulated objects significantly more than females, this manipulation was predominantly play-based (36). Conversely, females exhibited a more diverse array of early experiences with objects, such as breaking and plucking, which was more indicative of preparation for subsistence tool use. Thus, sex differences may arise from divergent experiences with objects during infancy. However, this finding does not explain the cause of variability in early object manipulation.

A 4-year longitudinal study on 14 infant-mother dyads in Gombe, Tanzania, assessed whether sex differences in the learning process precede the female bias in chimpanzee termite-fishing (5). The researchers found that female infants spent more time termite-fishing, peered at their mothers more, and successfully fished significantly earlier (ca. 27 months) than male infants. The congruence between the termite-fishing technique of female offspring and their mothers was greater than between male offspring and their mothers, which was interpreted to indicate that females learn the skill more via imitation (process-focused learning) and males more via emulation (product-focused learning).

These results suggest that the female bias in termite-fishing stems in part from divergent motivations to learn the skill during the acquisition period. However, no studies have assessed factors that might contribute to this variation or whether differences in peering exists in other technological behaviours, such as in stone tool use contexts. In turn, there has been no research into whether variation in peering, and potential differences in social learning mechanisms employed, produces variation in the long-term technological proficiency of individuals.

Because of the inherent role of model characteristics (in terms of, for example, tolerance) in the amount of time learners can peer, we argue that the proportion of complete nut-cracking sequences the learners peer at during their ELP is a more accurate measure to capture variation in learner peering than measuring total peering duration. Indeed, complete attention on the model performing the target behaviour is needed to ensure accurate observations (11). As such, the present study assessed the proportion of learner’s peering events that were directed at the entirety of the model’s nut-cracking bout, defined as the period (in seconds) of using a stone hammer to crack one nut on a stone anvil. Variation in model tolerance to peering and the number of learning opportunities provided (which impacts exposure to the behaviour) by the maternal model also greatly influence tool use proficiency (4). Tolerant models also afford more learning opportunities as they allow the learner to be proximate whilst peering at the skill being modelled, interact with the model and their tools to learn about the mechanics of the skill, and scrounge the extracted resource so that the behaviour is positively reinforced (4, 30, 37). Chimpanzee mothers in Gombe show the same amount of tolerance (defined as allowing their offspring to do whatever they were trying to do, such as peer or steal a tool) towards male and female infants learning to fish for termites (37). This research also observed “negative” behaviours mothers showed towards their infants (i.e., shoving, moving, and avoiding infants at termite mounds), but it did not assess whether differential exposure to this intolerance affected the learners’ behaviour. Given that intolerance is a form of punishment (thereby decreasing the likelihood of the behaviour being repeated) (38), it is a crucial variable to measure when assessing the ontogeny of technological efficiency.

Moreover, differences in social learning opportunities are thought to predict individual variation in tree-hole tool use in Sumatran orangutans (4). Data gathered over 3 years of the Bossou chimpanzees found that mothers differ in the proportion of time spent ant-dipping and hence provide varying degrees of learning opportunities for their infants (14). Infants exposed to higher rates of ant-dipping peering opportunities (having mothers who spent > 0.2% of their total activity time ant-dipping) started to ant- dip significantly earlier, dipped for longer, and made fewer dipping errors than offspring of low opportunity mothers (< 0.2% of total activity time ant-dipping) (14). For stone tool technology, the acquisition of nut-cracking in wild bearded capuchins is supported by tolerant models providing opportunities to observe (via stimulus enhancement) and practise the skill (39). Although this research did not assess variation during the developmental period, it highlights the importance of model opportunity provision during the acquisition of lithic technology.

In chimpanzees, the influence of maternal behaviour on nut-cracking outcomes have previously been investigated in 11 immature chimpanzees (aged 2.4–8 years) in the Taï National Park, Côte d’Ivoire (40). The 4-year study assessed “maternal scaffolding” (providing nut-cracking learning and practising opportunities for their offspring) and mother-initiated nut-sharing on: 1) the offspring’s task understanding (defined as using all three nut-cracking components), 2) the proportion of successful cracking bouts, and 3) efficiency (nuts per minute). The researchers found that predicted maternal scaffolding did not significantly affect infant nut-cracking outcomes, and although using the mother’s hammer did positively affect bout success and efficiency—likely by providing information on optimal hammer properties—offspring age had the largest influence on nut-cracking skills.

Although these findings appear to contradict the premise that opportunity provision and tolerance predict nut-cracking outcomes, the study did not encompass all the known elements of tolerant behaviour, such as allowing proximity and touch (30). Likewise, the research only investigated mothers’ “supportive behaviour” (scaffolding and nut-sharing) and did not assess *in*tolerance shown by the mothers (previously termed “negative” behaviours) (37). Intolerance is an important variable as it is an active model behaviour that directly affects the learner’s ability to be proximate to the behaviour and to scrounge extracted food items—factors known to be important for maintaining motivation during protracted learning periods (32). Conversely, tolerance to infant scrounging, proximity, and interaction is typically passive. Both tolerance and intolerance should be studied to understand the full suite of interactions learners experience with their selected models. Also, the research was conducted on sub- adult chimpanzees over a short period and so could not assess the critical longer-term influence of maternal behaviour on offspring nut-cracking efficiency.

To date, research on the ontogeny of tool use in wild primates has relied on relatively short-term observations (over a maximum of 4 years) and has often used cross-sectional data. However, given chimpanzees’ long lifespans and the protracted periods required to develop these skills (5, 8, 27), these data are insufficient for assessing the ontogeny of skill proficiency. The production of video archives by long-term field sites is now offering insights that would not be possible based on data from relatively short-term studies. For example, research using 25-years of footage (1992–2017) of the Bossou chimpanzees established stable and reliable individual-level variation across four measures of nut-cracking efficiency (time taken to crack one nut, number of strikes per nut, success rate, and number of times the nut was displaced from the anvil due to the hammer strike) in individuals aged 6+ (post- ELP) (9).

Here, we present a study aiming to establish the source of this variation by investigating the effect of experiences during the ELP (≤ 5 years old) on individuals’ long-term nut-cracking efficiency (≥ 6 years old). Four key aspects of nut-cracking skill development were recorded: 1) how frequently the learner peered at the model’s entire nut-cracking bout, 2) the amount of tolerance the models showed to the learner during peering, 3) the amount of *in*tolerance the models showed to the learner during peering, and 4) the social learning opportunities provided by the learner’s mother (see Table 1). Using 25 years of longitudinal data (1992–2017), this research tested whether variation in the ELP predicted the four measures of post-ELP efficiency of ten wild chimpanzees.

**Table 1.** Definitions of developmental measures.

| Measure | Definition |
| --- | --- |
| Whole bout peering | Where the whole of the model's nut-cracking bout was peered at, the peering event was recorded as "whole bout". Where only a segment of the bout was peered at, the peering event was recorded as "not whole bout". |
| Opportunity provision | The total proportion of time that the maternal model provided opportunities for their infant/juvenile to learn to crack nuts. This was a measure of the total amount of time the focal subject's mother was cracking nuts in the outdoor laboratory with the focal present, over the continuous period of when the focal subject's mother was on camera in the outdoor laboratory whilst the focal is also present (14). |
| Model tolerance | Where the model permitted the peering learner to scrounge the extracted kernel, interact with the model (e.g., touching the model's arm or body) or the model's tools, or be proximate ($\leq 1$ m away) to the model during the nut-cracking bout (28, 30). |
| Model intolerance | Where the model physically (e.g., shoving, hitting) or vocally (e.g., grunting, barking) rejected or resisted a peering learner's attempts to scrounge, interact with the model or the tools, or be proximate ($\leq 1$ m away) whilst they were engaged in a nut-cracking bout (37). |

## Results

### Variation in early learning period experience

Learning period data were collected for 13 individuals (six female and seven male, ages 0–5). Results show significant variation around the average for each of the four developmental measures across these 13 infants (see Figure 2 for observed *t*-values). For the ten individuals (five female and five male, ages 6–20) that we have data on their post-ELP, there was one who peered at the whole nut-cracking sequence significantly less than average (*n* = 1), and four who peered at the whole nut-cracking sequence significantly more than average (*n* = 4). Also, one learner received significantly fewer learning opportunities from their mother than average (*n* = 1), and two received significantly more than average (*n* = 2). For *model tolerance to peering*, no learners experienced significantly more tolerance than average but two experienced significantly below average tolerance (*n* = 2). For *model intolerance to peering*, four learners experienced significantly less intolerance than average (*n* = 4), and two experienced significantly more than average (*n* = 2).

**Figure 2.**
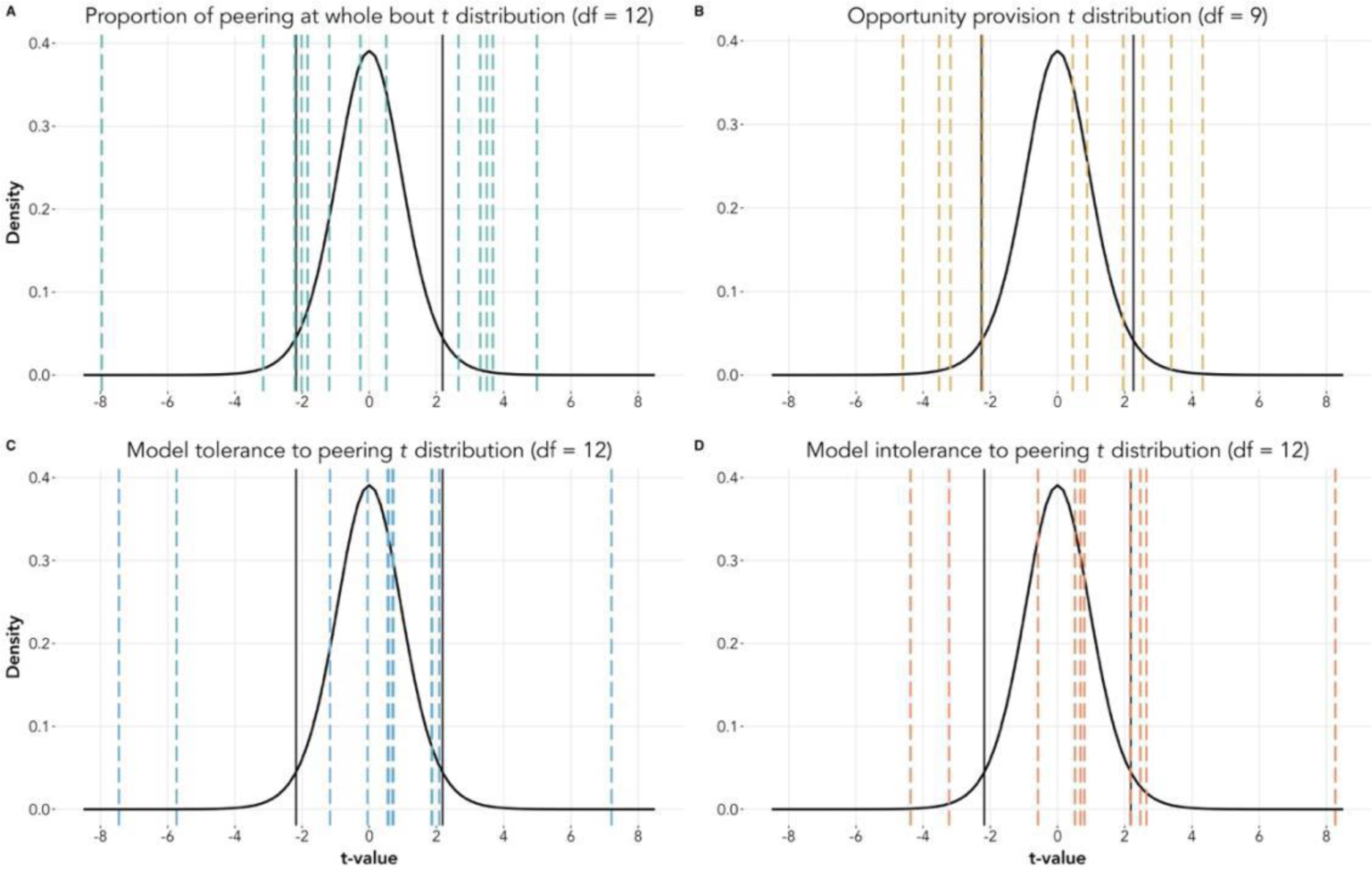
*t* distributions showing the variation in (A) proportion peering at the whole nut-cracking bout, (B) opportunity provision from the mother, (C) model tolerance to peering, and (D) model intolerance to peering during the development of nut-cracking. *Notes*: The black vertical lines indicate the 95% two-tailed critical *t-*values. The dashed vertical lines indicate the observed *t*-values for the focal subjects. Two chimpanzees shared the same *t*-score (-1.16) for model tolerance to peering, which explains why only 12 dashed lines exist in plot C. Five chimpanzees shared the same *t*-score (three for -4.36 and two for -0.56) for model intolerance to peering, which explains why only ten dashed lines exist in plot D. Observed *t*-values that fall between the black vertical lines show average scoring individuals. Observed *t*-values that fall beyond the black vertical lines are significantly different to average.

### Influence of the models on post-ELP nut-cracking efficiency

Exposure to social learning opportunities from the mother had a significant negative effect on post-ELP *strikes per nut* (number of strikes to crack one nut; *z* = -4.818, *p* < 0.001) but no effect on log *bout duration* (time taken to crack one nut; *t* = -1.946 *p* = 0.052), *success rate* (proportion of bouts that end in the whole kernel, the partial kernel, and no kernel being extracted; *z* = 1.453, *p* = 0.146) or *displacement rate* (number of times the hammer strike displaces the nut from the anvil; *z* = -0.907, *p* = 0.364).

Mean model tolerance had a significant negative effect on log *bout duration* (*t* = -2.395, *p* < 0.05) and *strikes per nut* (*z* = -2.978, *p* < 0.01), but no effect on *success rate* (*z* = -1.602, *p* = 0.109) or *displacement rate* (*z* = 0.505, *p* = 0.613). Mean model intolerance had a significant positive effect on log *bout duration* (*t* = 6.683, *p* < 0.001) and *strikes per nut* (*z* = 2.164, *p* < 0.05), but no significant effect on *success rate* (*z* = 0.229, *p* = 0.819) or *displacement rate* (*z* = 0.626, *p* = 0.613).

### Influence of the learner on post-ELP nut-cracking efficiency

The proportion of peering events in which the whole nut-cracking bout was observed had a significant positive effect on *strikes per nut* (*z* = 2.995, *p* < 0.01), and *displacement rate* (*z* = 5.185, *p* < 0.001), and a significant negative effect on *success rate* (*z* = -4.180, *p* < 0.001). There was no significant effect on log *bout duration* (*t* = 1.912, *p* = 0.056).

Post-ELP age had a significant negative effect on log *bout duration* (*t* = -4.514, *p* < 0.001), *strikes per nut* (*z* = -2.492, *p* < 0.05), and *displacement rate* (*z* = -6.537, *p* < 0.001), and a significant positive effect on *success rate* (*z* = 6.395, *p* < 0.001). There was also a significant effect of sex, with being male having a positive effect on *strikes per nut* (*z* = 2.985, *p* < 0.01), a significant negative effect on *success rate* (*z* = -2.420, *p* < 0.05), and no effect on *displacement rate* (*z* = 1.314, *p* = 0.189). Model coefficient estimates are shown in Figure 3.

**Figure 3.**
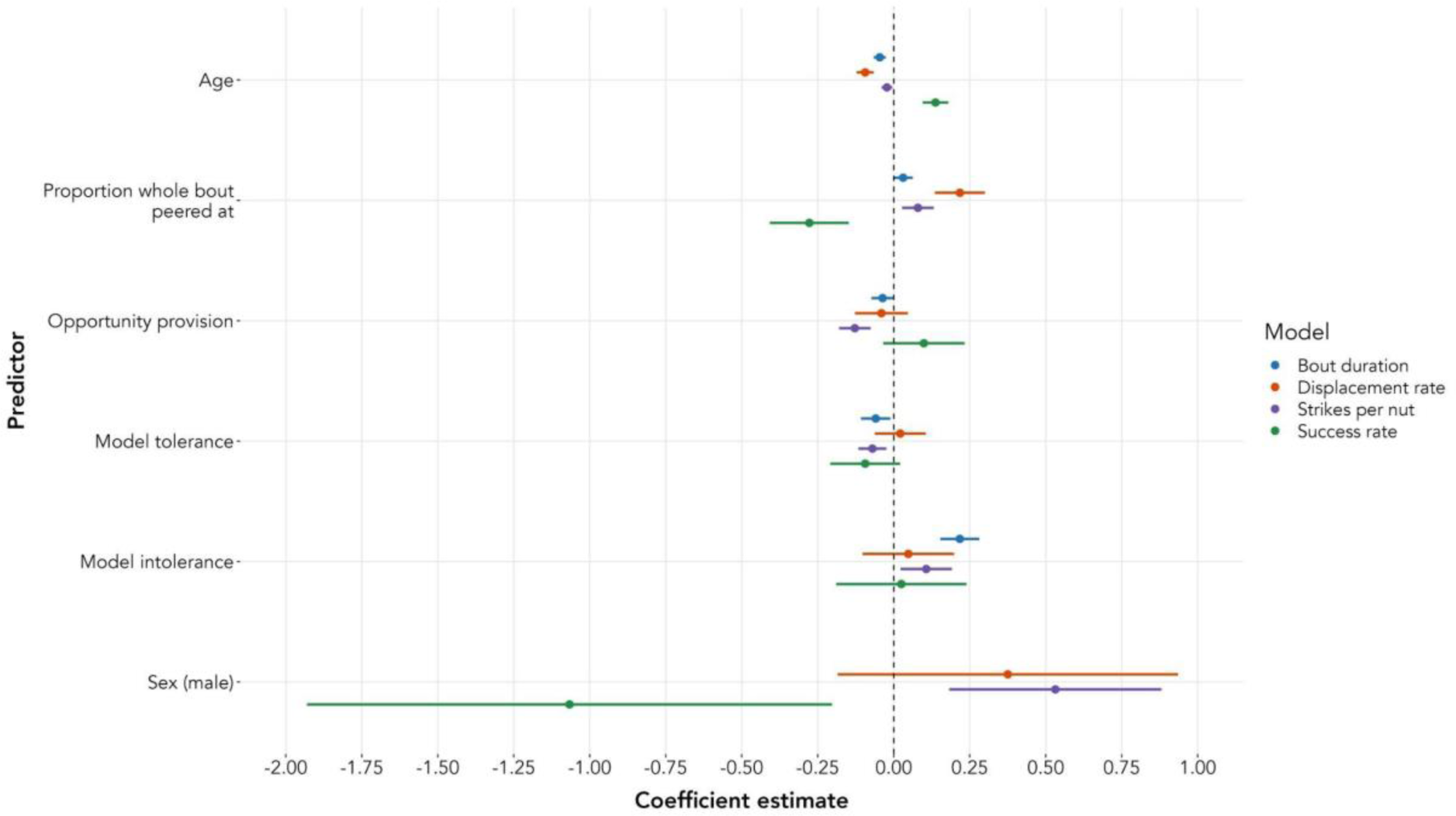
Coefficient estimates for each predictor variable from the model outputs of the four efficiency models. Dots indicate the estimate, and whiskers show the 95% confidence interval. The dashed vertical line at zero indicates no effect of the predictor on the outcome.

## Discussion

Understanding the factors driving variation in chimpanzee tool use is critical for understanding the selective pressures on primate and early hominin cognitive and technological evolution. This longitudinal study shows that divergent experiences during the development of tool use are related to variation in the subsequent technological efficiency of wild primates. Specifically, we show that variation in the opportunities to learn socially, as well as the amount of tolerance and intolerance experienced whilst attempting to peer at nut-cracking models during the learning period, are associated with the individual’s post-ELP nut-cracking efficiency (9). This has implications for theories of the origins of tool use (17), particularly as we found that model *in*tolerance was a stronger predictor than model tolerance for subsequent skill efficiency (in terms of bout duration).

Our results offer longitudinal support for the hypothesis that the number of social learning opportunities and the amount of tolerance to peering (i.e., being allowed to scrounge, interact with, and be in close proximity to nut-cracking models) provided during development predicts future technological proficiency (4). Individuals exposed to more social learning opportunities from their mother and those who experienced more tolerance when attempting to peer subsequently required fewer strikes to extract the kernels (which reflects energy spent relative to energy gained) (9). Additionally, learners who were exposed to more tolerance from models subsequently had shorter nut-cracking bouts—an indicator of energy intake rate (9). Being exposed to social learning opportunities by tolerant models (including local and stimulus enhancement and observational learning) is considered to be a form of niche construction (32), which is facilitated by the enduring nature of stone tools (41). The distinctive sound of the nut-cracking percussion (42) along with the capacity for young learners to reuse adults’ tools provides both the physical affordance to use the tool composite and the motivation to do so (41).

Conversely, exposure to social learning opportunities presented by mothers and model tolerance to peering did not predict subsequent success rate or displacement rate, which are indicators of strike and kinetic energy accuracy (9, 43, 44). Similarly, maternal opportunity provision did not predict subsequent log bout duration. It could be that these components of nut-cracking efficiency are honed and impacted by different developmental processes, such as learning by trial-and-error, or by genetic processes relating to strength and coordination. Alternatively, the results may reflect the fact that our research only focussed on the social learning opportunities provided by mothers, meaning that all learning opportunities available to the chimpanzees during the development of nut-cracking were not captured. Nonetheless, our results provide an important step in highlighting the long-term consequences of differential exposure to learning opportunities on skill efficiency.

Our findings that learners who experience more intolerance (e.g., being shoved away or not being allowed to scrounge) whilst attempting to peer at nut-cracking models were less efficient (in terms of bout duration and strikes per nut) later in life advances van Schaik and colleagues’ theory of the ontogeny of primate technological behaviour (4). Intolerant behaviour punishes the learners for trying to observe the skill, meaning that they are unable to watch the behaviour at close range to learn the technique or gain motivating positive reinforcement from scrounging. Again, the future success rate and displacement rate of the individual was not impacted by intolerance to peering, further suggesting that these aspects of efficiency develop via different learning processes.

This supports the idea that individual variation can arise from “individual learning of social learning” (p. 215), whereby the rate of learning and the learning strategies used are phenotypically plastic, contingent upon previous social learning interactions (13). Experiencing high degrees of intolerance whilst attempting to learn by observation (i.e., peering) may force learners to rely upon other forms of learning (whether social or asocial). These learning mechanisms may be suboptimal for acquiring complex extractive foraging skills and becoming technologically efficient, hence the resultant reduction in relative efficiency post-ELP for these individuals. Future research investigating whether the rate of individual and social learning of the skill varies depending on the amount of intolerance individuals experience from models would shed further light on this process.

Regarding the influence of the learner’s behaviour on nut-cracking efficiency, we found that, counter to predictions, learners who peered at a higher proportion of whole nut-cracking bouts subsequently became *less* efficient nutcrackers (in terms of more strikes per nut, reduced success rate, and more nut displacements). Although not statistically significant, these learners also tended to have longer nut-cracking bouts. It is possible that the “whole-bout-peerers” could have peered less overall, meaning that although they observed a larger proportion of whole bouts, they also observed fewer bouts overall relative to their conspecifics. However, there was only a weak correlation (*r* = 0.26) between number of peering events and proportion of whole bout peering. Alternatively, the finding that “whole-bout-peerers” became less efficient nutcrackers may be an artefact of the binary nature of the “whole bout peered at” measure used in this research, with a peering event being coded as either a “whole bout” or “not the whole bout”. Notwithstanding this possibility, the fact that the proportion of whole nut-cracking bouts peered at during the learning period predicted three of the four measures of efficiency highlights the importance of this variable in the ontogeny of chimpanzee technological efficiency.

We tentatively suggest that these results may reflect the process of cultural transmission: peering at the whole bout is not to learn how to perform the behaviour efficiently, but rather to learn ‘the way we do things’ as a group (45, p. 433). Therefore, the learners may become less efficient because they are replicating superfluous cultural components that do not improve skill efficiency. Peering at the whole nut-cracking sequence of models increases the likelihood of the cultural behaviour being maintained, as the technique can be more faithfully transmitted (11). We found that the majority of all 436 peering events (60.55%) involved the learner peering at the whole of the model’s nut-cracking bout, with the proportion increasing as infants aged (see Figure S1). Although the increase in the proportion of whole bout peering over the course of the ELP is likely to reflect, at least in part, developmental changes in the ability to sustain attention, it may also illustrate this process of cultural transmission and maintenance. Research is now needed to determine the degree of congruence between the technique of the models whose whole bouts were peered at, and the learners’ subsequent nut-cracking techniques (as was previously done in termite-fishing in Gombe) (5). This will shed light on whether learners who peer at the whole nut-cracking bout more are imitating the model’s behaviour, rather than emulating the sequence to achieve the same outcome, thereby advancing understandings of the social learning mechanisms employed by wild chimpanzees (46, 47). At the very least, the finding reiterates the need to test assumptions of the social learning capabilities of chimpanzees in wild populations learning naturally occurring skills, rather than in artificial tasks in captive individuals (48).

This research assessed the combined influence of the learner and model on the long-term technological efficiency of primates. We show the importance of both the learners’ and the models’ (maternal and non-maternal) behaviour on subsequent nut-cracking efficiency. Although our sample is limited to the relatively small size of the Bossou community (which is in the lower end of the normal range for chimpanzee populations) (49), this population, currently very close to extinction, is the only one where long-term stone tool use has been recorded yearly using a systematic video recording approach. It thus offers a unique and unrepeatable opportunity to disentangle the relative contribution of both the “master” and the “apprentice” during the learning period. Observational learning plays an integral role in the acquisition of nut-cracking by wild chimpanzees. Follow up research will ascertain whether any model-based social learning strategies (50) are involved in non-maternal model choice, such as kin or proficiency biases (51, 52).

Age had a significant effect on all post-ELP efficiency measures. As chimpanzees got older, their bout durations decreased, they required fewer strikes to extract the kernel, their success rate increased (i.e., their bouts were more likely to end with the whole kernel being extracted), and they made fewer displacements per nut. We also found a significant effect of sex for two measures of efficiency, with male chimpanzees requiring more strikes per nut and having a lower success rate than females. However, these results should be viewed with caution as previous research found there was no effect of sex in chimpanzee nut-cracking efficiency across a larger sample and dataset (9). Results may be a function of the sample size, as data on the learning and post-ELP across the 25-year period sampled existed for ten chimpanzees, compared to the 21 individuals in the previous research (9). Alternatively, it could be the case that sex differences exist earlier in life (the maximum age in this data set was 20 years old), and that these disappear later in life in Bossou.

This research has two main limitations. First, as mentioned, the binary measure for whether the individual peered at the whole nut-cracking bout or not may have been too coarse to sufficiently capture variation in the learners’ peering behaviour, and its effect on subsequent efficiency. Future research should investigate the exact proportions of the models’ bouts the learners peered at, and the reasons for looking away (e.g., to manipulate objects, because of a loud noise, to engage in a social interaction, etc.) to better understand the role of peering at the entire behavioural sequence on the acquisition of the skill.

Second, the research was inherently constrained by using pre-collected and historic data. At times, events of interest (particularly peering) were happening on camera but the camera angle or video quality hindered data coding. While we doubt that this biased our results, it did reduce the size of the overall data set for analysis. Moreover, the observational nature of the research limits our ability to establish a causal link between experiences during the ELP and subsequent efficiency. However, these constraints are outweighed by the advantages of using long-term video records, which allow tests of questions that are not possible using short-term field research alone (53).

We provide critical longitudinal data strongly suggesting that observational social learning processes are fundamental to the acquisition of nut-cracking. This has important implications for the use of animal culture in conservation. Nut-cracking by West African chimpanzees is protected by the United Nations’ Convention on the Conservation of Migratory Species of Wild Animals and is recognised as being vital for the conservation of the subspecies (54). Our data reiterate the warnings of the disturbance hypothesis: where the core components of social transmission are impacted by human pressures, such as hunting and habitat degradation and fragmentation, behavioural repertoires and fitness-enhancing skills and knowledge will be severely disrupted (55). Long-term data, such as those presented here, can contribute significantly to understanding whether culture can and should be used in conservation policy. It is clear that if nut-cracking is to be protected, the social learning processes through which the behaviour is acquired must be too. Ultimately, this means protecting the chimpanzees.

## Materials and Methods

### Subjects and Data Acquisition

#### Study site

The wild chimpanzees neighbouring Bossou village in the tropical wet seasonal climate of south-eastern Guinea (7°38’71.7’N, 8°29’38.9’W) have a home range of 15km^2^ and a core area of 7km^2^ within primary and secondary forest (56). The community has been under investigation since 1976, with the population size always being relatively small (9–21 individuals during the research period of 1992– 2017) (57, 58).

#### Study materials

The nut-cracking behaviour of the Bossou chimpanzees has been recorded in every dry season (December–February) since 1988, with videos taken from behind a grass screen from at least two standardised positions (59). This has produced a long-term video archive from within the so-called “outdoor laboratory”: a 7m x 12m site on Mont Gban (7°38’41.5’’ N, 8°29’50.0’’ W) where locally sourced stones and nuts are experimentally provisioned (8, 57). This study used the 1992–2017 footage, excluding 2001 and 2011 as footage was not taken in these years. The 2009 footage was also excluded, as only experimentally introduced *Coula edulis* nut-cracking bouts were peered at that year.

#### Measures

This study assessed four developmental measures (see Table 1): proportion of peering at the full length of the models’ nut-cracking bouts, exposure to social learning opportunities by the mother, tolerance experienced from all models whilst peering, and intolerance experienced from all models whilst attempting to peer. Peering is ‘the attentive and sustained close-range watching of the activities of a conspecific’ (34, p. 5). Here, a peering event was defined as the period (in seconds) where the learner’s gaze was directly oriented towards a model’s hand, arm, nut, or stone tool composite during a nut-cracking bout (8). A nut-cracking bout was defined as the length of time between the chimpanzee lifting the hammer for the first strike and the hammer making contact for the final time (9, 24). Each peering event occurred at the bout-level, meaning that a new peering event was recorded for each new oil palm nut the model cracked. The name of the model and their relation to the learner was recorded for each peering event. Whether or not the learner peered at the whole bout was recorded. Whether the model permitted learner scrounging, proximity, or interaction whilst peering (tolerance), or expressed physical or vocal intolerance towards the learner attempting to peer, was also recorded for every peering event.

The four separate (but stable and reliable) measures of post-ELP efficiency were used: the amount of time taken in seconds to crack one nut (“bout duration”), the number of strikes required to crack one nut (“strikes per nut”), the “success rate” of the individual, and the number of times the nut was displaced from the anvil due to the hammer strike (“displacement rate”) (9).

#### Data collection

Data for the four developmental measures were collected using *Behavioural Observation Research Interactive Software* (BORIS; v. 7.13.9) (60). Data on post-ELP efficiency were collected as part of previous research (9). All-occurrence sampling was employed to gather data from all visible peering events during the video archive for each focal individual for every available year of their learning period (six years per individual from 0–5 years old, inclusive). Visible peering events were those where the focal was facing the camera or where they were not directly facing the camera, but their gaze was observable. For model tolerance to peering and model intolerance to peering, data were gathered from all peering events using the “modifier” function in BORIS. All instances of tolerance and intolerance the model showed to the learner during a peering (or attempted peering) event were recorded. The ethogram and modifiers used during data collection can be found in Tables S1 and S2. For opportunity provision, all-occurrence sampling was used to gather data on the proportion of the total observation time that the learners’ mothers were nut-cracking in the outdoor laboratory. Data were not obtained from 1994 or 1995, as rapid and shaky camera movements hindered the ability to accurately measure the length of nut-cracking.

Since 1976, there have been 28 mother-infant dyads (see Table S3) (61), but footage of the full nut-cracking learning period is only available for 19 of these infants. Of these 19 chimpanzees, one was born after the study period, and one’s learning period commenced prior to 1992 and so they were excluded from the study. Four chimpanzees died before age three, and so not enough footage existed for them to be included. A further three chimpanzees either died before age six or reached age six after 2017, and so there is no data on their post-ELP efficiency. As such, developmental data were collected from 13 infants, and the role of the developmental period on post-ELP efficiency was assessed in ten of these individuals. Three of these infants (Nto, Poni, and Peley) had mothers who did not crack nuts, and post-ELP data exist for all three.

Details on the data collection protocol for the post-ELP efficiency of the Bossou chimpanzees has been previously published (9). In short, data were gathered from at least 20 bouts (where possible) for each year the ten focal subjects (aged 6 ≤) were visibly cracking oil palm nuts in the footage from 1998– 2017.

For the developmental period, a total of 436 peering events, 466 instances of model tolerance, and 52 instances of model intolerance were recorded across 13 chimpanzees (six female and seven male, ages 0–5) from over 680 hours of footage from 1992–2017 (see Table S4). Of the 436 peering events, 264 (60.55%) involved the learner peering at the model’s entire nut-cracking bout. For the post-ELP efficiency measures, a total of 1098 nut-cracking bouts were recorded across the ten focal chimpanzees (five female and five male, ages 6–20) from 1998–2017 (9).

#### Ethics

During the 1992–2017 research period, research authorisation and ethical approval were collectively given to TM, the leader of the Bossou Project of Kyoto University, from Guinean authorities (Direction Générale de la Recherche Scientifique et de l’Innovation Technologique, DGERSIT, and the Institut de Recherche Environnementale de Bossou, IREB) based on the Memorandum of Understanding. Since data collected were all archival and required no human participants, no further ethical approval was required by the School of Anthropology and Museum Ethnography, University of Oxford, or the Social Science Division, University of Oxford.

### Statistical analysis

All analyses were performed using R (v. 4.3.2) (62) and RStudio (v. 2023.09.1+494) (63) for MacOS. Given the small sample size, variation in the four developmental measures was assessed by comparing observed *t*-values to critical *t-*values (*p* < 0.05, two-tailed). All observed *t-*values less than the lower critical *t*-value threshold were significantly below average (e.g., significantly fewer learning opportunities provided by their mother, etc.), and all observed *t*-values greater than the upper critical *t*-value threshold were significantly above average (e.g., significantly more learning opportunities provided by their mother). These *t*-values were then used as predictors in the regression models. As *t*-scores use the average of the data points for each case, this approach resolves the issue of the data often being sequential, which leads to non-independence due to temporal autocorrelation (64).

For bout duration and strikes per nut efficiency measures, all “failed” bouts (*n* = 146) were removed to ensure the analyses assessed the amount of time and energy exerted to retrieve the energetic reward (9). This left 952 post-ELP bouts for analysis. All bouts (*n* = 1,098) were analysed for success rate and displacement rate.

Mixed-effect models (with subject and mother as random intercepts) were constructed for each of the four efficiency outcome variables, with age, sex, whole bout peering proportion *t*-score, opportunity provision *t*-score, model tolerance *t*-score, and model intolerance *t*-score as predictors.

The bout duration data were log transformed to normalise the positively skewed distribution. The linear mixed-effects model for log bout duration was fitted using the *lme4* package (65). The sex predictor had an issue of multicollinearity (VIF = 7.07), and so was removed from the model. A zero-truncated Poisson generalised linear mixed model (GLMM) was fitted for strikes per nut using the *glmmTMB* package (66). Overdispersion was detected, so a zero-truncated negative binomial GLMM with quadratic parameterisation was fitted to counteract this issue. For success rate, a cumulative link mixed model was fitted using the *ordinal* package (67) as the outcome variable data were ordered in order of increasing efficiency (Failed, Smash, Successful) (44). Given the substantial number of zeros in the displacement rate data (*n* = 754), a zero-inflated (zi ∼ 1) Poisson GLMM was fitted using the *glmmTMB* package. This model was over-dispersed, so a zero-inflated (zi ∼ 1) negative binomial GLMM was fitted to the data. However, there was a model convergence problem, likely due to over-parameterisation with a small sample size. Removing the random intercepts for subject and mother fixed this issue; the final model thus did not contain random intercepts for subject and mother.

Allowing the intercept to vary by subject did not significantly improve the model fit for any of the models, as indicated by larger AIC scores. The assumptions of all models were checked *post hoc* (see Supporting Information).

Independent, hypothesis-blind coders reviewed 14.5 hours of footage for inter-coder reliability analyses, which totalled 81 peering events. Unweighted Cohen’s κ indicated substantial agreement between coders for model tolerance (κ = 0.797, *z* = 10.7, *p* < 0.001), model intolerance (κ = 0.741, *z* = 6.7, *p* < 0.001), and whether the whole bout was peered at or not (κ = 0.833, *z* = 7.51, *p* < 0.001). For opportunity provision, two-way, single-rating random-effects intraclass correlation (ICC) analyses were performed using the *irr* package (68). Results indicated excellent consistency between coders for both the start (ICC(C,1) = 1, F(26,26) = 29775, 1 < ICC < 1, *p* < 0.001) and stop (ICC(C,1) = 1, F(26,26) = 5403, 0.999 < ICC < 1, *p* < 0.001) time of maternal opportunity provision.

### Code availability

All analysis code needed to reproduce the results and figures reported in the paper and the Supporting Information can be found in the following public repository: https://github.com/arranjdavis/chimpanzee_nut_cracking_ontogeny

## Acknowledgements

The video data for this research was provided by Tetsuro Matsuzawa and the Primate Research Institute, Kyoto University, Japan. The original video archive was digitised, organised, and systematised by Daniel Schofield, Department of Engineering Science, University of Oxford. Special thanks to all the field assistants and researchers at Bossou, specifically Henry Didier Camara, Cé Vincent Mamy, Gouanou Zogbila, Boniface Zogbila, Jules Doré, Gouanou Goumy, Tino Camara, Pascal Goumy, Paquílé Cherif, Jiles Gondo Doré, and Marcel Doré. Thank you to Alexander Mielke for help cataloguing the Bossou video archive and for early discussions about this research, and to Dora Biro and Kathelijne Koops for feedback. Thank you to Rebecca Funnell and Carolyn Gover for research assistance with hypothesis-blind coding for the inter-rater reliability analyses.

