## Supporting Information for "The ontogeny of chimpanzee technological efficiency"

*Corresponding author

### Supporting Information Text

#### Statistical Analysis

***Assumption checks***. *Post hoc* analyses were run to check the model assumptions. For the *bout duration* linear model, multicollinearity between the predictor variables was checked using the *vif* function from the *car* package (1). The VIFs did not exceed 3.34, indicating no issues with multicollinearity (2). The normality of the residuals was checked visually using a histogram and QQ-plot and homoscedasticity was assessed with a plot of fitted values against squared residuals. Influential observations ($n$ = 65) were identified using the *cooks.distance* function from the *car* package. Most of the influential observations were either less than a second in duration or more than 15 seconds in duration (i.e., over three times the average bout duration). We nonetheless decided to retain these data in the model because they represent the variation that we are interested in.

For the *success rate* cumulative link model, the normality of the surrogate residuals (3) was checked using a QQ-plot and the homoscedasticity was checked with a plot of fitted values against surrogate residuals (4) using the *sure* package (5). For the *strikes per nut* and *displacement rate* glmmTMB models, the normality of the simulated residuals and homoscedasticity were checked using the *DHARMa* package (6). No significant problems of homoscedasticity were detected for either model when the model residuals were plotted against the predicted values. The QQ-plot of expected versus observed residuals indicated the simulated residuals did not deviate from normality for *displacement rate*. However, the simulated residuals deviated from normality for *strikes per nut*, likely due to the presence of nine outliers (indicated by a two-sided DHARMa bootstrapped outlier test, $p < 0.001$). Nearly all the outliers were strike counts that were over three times the average (over 12 strikes per nut, maximum of 20 strikes), but we again decided not to remove these observations as they represent the variation that we are interested in.

***Table S1.*** Ethogram of behaviours coded in BORIS.

| **Behavioural Category** | **Type** | **Definition** |
|---|---|---|
| Start Peering | Point event | The start of the period when the focal subject's gaze is directly oriented towards a model's hand, arm, nut, or stone tool composite during a nut-cracking bout. |
| End Peering | Point event | When the focal subject's gaze is no longer oriented towards the model's hand or arm or towards the nut and stone tool composite. |
| Start Opportunity | Point event | When the focal's mother commences nut-cracking whilst the focal subject is in the outdoor laboratory. |
| End Opportunity | Point event | The focal's mother ceases to crack nuts whilst the focal subject is in the outdoor laboratory. |
| Total Visibility | State event | The continuous period of when the focal subject's mother is on camera whilst the focal subject is in the outdoor laboratory. |

***Table S2.*** Modifiers applied to the ethogram in BORIS.

| Behaviour | Modifier | Definition |
|---|---|---|
| Start Peering | Model name | The name of the individual the focal subject was peering at during their nut-cracking bout. |
| | Maternal model | Whether the model was the learner's mother or not. |
| End Peering | Complete peering event | Whether or not the full peering event was able to be recorded because of the video footage itself. |
| | Whole bout | Whether or not the learner was peering for the whole nut-cracking bout. |
| | Model name | The name of the individual the focal subject was peering at during their nut-cracking bout. |
| | Maternal model | Whether or not the model was the learner's mother. |
| | Model tolerance | Whether the model permitted any of the following learner behaviours: scrounge, tool interaction, model interaction, proximity. |
| | Model intolerance | Whether the model expressed physical or vocal intolerance towards the learner. |
| End Opportunity | Complete opportunity | Whether or not the full opportunity was able to be recorded because of the video footage itself. |
| | Learner behaviour | Which behaviour(s) the infant was engaged in when their mother was cracking nuts: solitary play, social play, object play, behavioural episode, bout, feeding, drinking, resting, peering, clinging, unknown. |
| | Scrounge events | How many times the focal subject took the kernel extracted by the maternal model to eat themselves. Only record these scrounge events where the focal subject took the kernel but did not peer at the nut-cracking bout. |

***Table S3.*** Genealogy of the Bossou chimpanzees since the formation of the field site in 1976.

| Mother | Born | Disappeared | Offspring | Offspring sex | Born | Disappeared |
|---|---|---|---|---|---|---|
| Jire | ~1958 | Present | Ja | Female | 1983 | Feb 1993 |
| | | | Jokuro | Female | 1989 | Jan 1992 |
| | | | Juru | Female | 1993 | Dec 2001 |
| | | | Jeje | Male | 1997 | Present |
| | | | Jimato | Male | 2002 | Dec 2003 |
| | | | Joya | Female | 2004 | 2013 |
| | | | Jodoamon | Female | 2009 | 2010 |
| Yo | ~1961 | 2022 | Yunro† | Female | 1984 | Feb 1993 |
| | | | Yolo | Male | 1991 | Nov 2009 |
| Fana | ~1956 | Sept 2022 | Foaf | Male | 1980 | 2023 |
| | | | Fotaiu | Female | 1991 | Nov 2004 |
| | | | Fanle | Female | 1997 | Feb 2024 |
| Fanle | 1997 | Feb 2024 | Flanle | Male | 2007 | Aug 2016* |
| | | | Fanwa | Male | 2012 | Present |
| | | | Fangasi | Female | 2020 | 2021 |
| Fotaiu | 1991 | Nov 2004 | Fokaiye | Male | 2001 | Nov 2004 |
| Pama† | ~1967 | Sept 2013* | Pili | Female | 1987 | May 2001 |
| | | | Poni | Male | 1993 | Dec 2003 |
| | | | Peley | Male | 1998 | Sept 2013* |
| Pili | 1987 | May 2001 | Pokuru | Male | 1996 | May 2001 |
| Velu | ~1959 | Mar 2017* | Vube | Female | 1982 | Mar 1990 |
| | | | Vui | Male | 1986 | Jul 1999 |
| | | | Vuavua | Female | 1991 | Mar 2004 |
| Vuavua | 1991 | Mar 2004 | Veve | Female | 2001 | Dec 2003 |
| Nina† | ~1954 | Nov 2003 | Na | Male | 1985 | May 1996 |
| | | | Nto | Female | 1993 | Dec 2001 |
| Kai | 1950 | Dec 2003 | Kie | Female | 1976 | Dec 2003 |
| Kie | 1976 | Dec 2003 | Kakuru | Female | 1986 | Mar 1991 |

*Notes*: Data from "*The Chimpanzees of Bossou and Nimba,*" eds. T., Matsuzawa, T., Humle, & Y., Sugiyama, 2011, Appendix A, p. 404–5. * = Data from "*The chimpanzees in Bossou,*" https://www.greencorridor.info/en/chimp/profile/relationship.html. † = Could not crack nuts.

**Table S4.** Focal subject information and data summary.

| Subject | Sex | Learning period | Post-learning period | Nut-cracking mother | Age range | Proportion peering whole bout | Opportunity provision | Mean tolerance | Mean intolerance | Post-learning period bouts |
|---|---|---|---|---|---|---|---|---|---|---|
| Fotaiu | F | 1992–1997 | 1998–2003 | Yes | 0–11 | 0.606 | 0.675 | 0.758 | 0.030 | 90 |
| Vuavua | F | 1992–1997 | 1998–2004 | Yes | 0–12 | 0.500 | 0.403 | 1.100 | 0.100 | 106 |
| Yolo | M | 1992–1997 | 1998–2009 | Yes | 0–17 | 0.440 | 0.593 | 1.160 | 0.100 | 208 |
| Juru† | F | 1994–1999 | - | Yes | 0–5 | 0.750 | 0.613 | 1.090 | 0.136 | - |
| Nto | F | 1994–1999 | 2000 | No | 0–6 | 0.529 | - | 1.060 | 0.000 | 11 |
| Poni | M | 1994–1999 | 2000–2002 | No | 0–9 | 0.571 | - | 1.000 | 0.000 | 41 |
| Pokuru* | M | 1997–2000 | - | Yes | 0–4 | 0.222 | 0.366 | 1.440 | 0.333 | - |
| Fanle | F | 1998–2003 | 2004–2017 | Yes | 0–20 | 0.742 | 0.541 | 1.160 | 0.129 | 234 |
| Jeje | M | 1998–2003 | 2004–2017 | Yes | 0–20 | 0.704 | 0.642 | 1.090 | 0.185 | 236 |
| Peley | M | 1999–2004 | 2005–2012 | No | 0–14 | 0.733 | - | 1.000 | 0.133 | 133 |
| Joya | F | 2004–2010 | 2010–2012 | Yes | 0–8 | 0.810 | 0.556 | 0.667 | 0.000 | 25 |
| Flanle | M | 2008–2013 | 2014 | Yes | 0–6 | 0.492 | 0.448 | 1.010 | 0.129 | 14 |
| Fanwa† | M | 2012–2017 | - | Yes | 0–5 | 0.483 | 0.415 | 1.170 | 0.172 | - |

*Notes:* * = Died before the end of their learning period. † = No data on post-learning period nut-cracking efficiency.

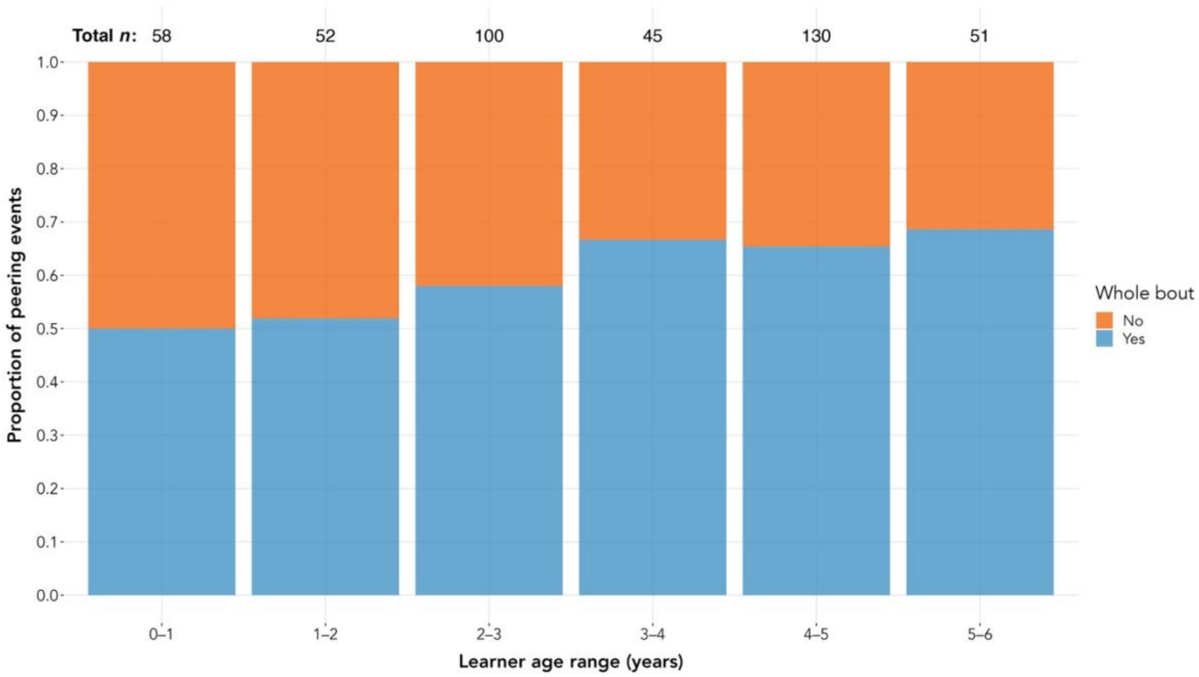


***Figure S1.*** Proportion of peering at the whole nut-cracking bout according to the learner’s age.

*Note*: The age range for each bar is inclusive of the lower value and exclusive of the upper value (e.g., $0 \leq x < 1$).
